# HDAC inhibitors rescue MeCP2^T158M^ speckles in a high content screen

**DOI:** 10.1101/2023.11.02.565272

**Authors:** Rodrigo Lata, Liesbeth Steegmans, Ranie Kellens, Marnik Nijs, Hugo Klaassen, Matthias Versele, Frauke Christ, Zeger Debyser

## Abstract

Rett syndrome (OMIM 312750) is a rare neurodevelopmental disorder caused by *de novo* mutations in the Methyl-CpG Binding Protein 2 (MeCP2) gene located on the X-Chromosome, typically affecting girls. Currently, available therapy for Rett Syndrome is only symptomatic. Rett syndrome symptoms first appear between 6 to 18 months of age, characterized by microcephaly and lack of motor coordination being the most prevalent. The disease continues to progress until adulthood when it reaches a stationary phase. More than 800 different mutations causing Rett syndrome have been described, yet the most common is T158M (9% prevalence), located in the Methyl-Binding domain (MBD) of MeCP2. Due to its importance for DNA binding through recognition of methylated CpG, mutations in the MBD have a significant impact on the stability and function of MeCP2. MeCP2 is a nuclear protein and accumulates in liquid-liquid phase condensates visualized as speckles in NIH3T3 by microscopy. We developed a high content phenotypic assay, detecting fluorescent MeCP2 speckles in NIH3T3 cells. The assay allows to identify small molecules that stabilize MeCP2-T158M and phenotypically rescue speckle formation. To validate the assay, a collection of 3572 drugs was screened, including FDA-approved drugs, compounds in clinical trials and biologically annotated tool compounds. 18 hits were identified showing at least 25% of rescue of speckles in the mutant cell line while not affecting wild-type MeCP2 speckles. Primary hits were confirmed in a dose response assay and in a thermal shift assay with recombinant MeCP2. One class of identified hits represents histone deacetylase inhibitors (HDACis) showing 25% speckle rescue of mutant MeCP2 without toxicity. This screening strategycan be expanded to additional compound libraries and support novel drug discovery.

## Introduction

Rett syndrome (RTT) is a rare X-linked neurological disorder affecting 1 in 15000 girls at birth. The disease is caused by *de novo* mutations in the methyl-CpG-binding protein 2 (*MECP2)* gene^1^. Generally, Rett syndrome patients show symptoms from the age of 6 to 18 months, losing acquired motor coordination or speech ability and displaying microcephaly^2^. Following the appearance of the first symptoms, the disease progresses rapidly until it reaches a plateau in most patients, while some patients will progress further until they lose mobility^3^. Since the severity of the disease is linked to the type of mutation and to the impact of mosaicism, Rett syndrome presents with a wide spectrum of symptoms and severity^1^. Indeed, more than 800 different mutations in MeCP2 have been described^4^. The majority are located in two functional domains:

(1) the methyl binding domain (MBD, 85 aa) and (2) the transcriptional repression domain (TRD, 104 aa)^4^. MeCP2 is expressed in two different isoforms, MeCP2E1 and MeCP2E2, which differ in function and expression patterns. MeCP2E1 has an additional 12aa, characterized by a seven alanine repeat followed by six glycines, that impact the protein turnover ^5^. MeCP2E1, the most prevalent isoform in neurons, shows reduced DNA-binding affinity and structural stability. The only structural difference between both isoforms is located in the NTD, which contributes to enhancing MBD’s affinity for DNA ^6,7^. Moreover, MeCP2E1’s expression levels vary in accordance with the circadian rhythm, while MeCP2E2 has a stable expression throughout the day ^5,8,9^. While the diverging expression patterns suggest distinct roles for MeCP2E1 and MeCPE2, the similar structure imply the potential for overlapping functions.^5^ Besides MBD, MeCP2 is intrinsically disordered, when associated with DNA. It forms droplets inside the nucleus due to liquid-phase separation. These droplets, which are functional, play an important role in MeCP2-chromatin interaction, affecting the condensation of the heterochromatin^10,11^. RTT-causing mutations located in the MBD affect chromatin compaction, impair DNA binding and thus the recruitment of MeCP2 binding partners such as HDACs, responsible for chromatin compaction^12^. It is known that the chromatin is organized differently in mouse and human cells. In the mouse nucleus, the chromatin forms clear pericentromeric domains were MeCP2 binds while in human cells, these pericentromeric domains are less prominent^13^.

MeCP2 is essential for arborisation and synapse density of nerve cells^14,15^. MeCP2E1 expression levels are low during early stages of development but increase at the start of neuronal differentiation^8^. In adults, the protein is expressed at high levels in the cortical regions, hippocampus, cerebellum, and olfactory bulb^13,16^.

The MBD of MeCP2 recognizes the CpG-islands in the promoter region of specific genes and the repressive function of MeCP2 is due to the transcription repression domain (TRD) that recruits several binding partners such as Sin3a and Histone deacetylases (HDACs)^17^.The main role of HDACs is to remove acetyl groups from histones, increasing gene repression^18^. Studies show that HDACs, particularly HDAC6, can regulate brain-derived neurotrophic factor (BDNF). BDNF plays an important role in neuronal plasticity and survival, specifically in the hippocampus. It is known that MeCP2 mutations affect BDNF levels in mouse models, however the role of BDNF in Rett syndrome is still unclear.

The most frequent missense mutation, present in approximately 10% of all Rett patients is T158M^4^. Threonine 158 plays an important role in the tandem Asx-Serine Threonine motif of the MBD of MeCP2 and is essential for binding to DNA containing 5-methylcytosine (5mC-DNA). The mutant protein shows reduced stability compared to the wild-type contributing to the loss of function^19,20^. The lower affinity and stability of MeCP2^T158M^ affect the localization of the protein. While the wild-type protein presents normal binding and clustering in NIH3T3 cell lines, this specific mutant presents impaired clustering as previously described^21,22^. The T158M mutation displays a mild to severe clinical score in most cases yet causes the most severe respiratory problems,^23^ which represent a serious complication in patients leading to frequent respiratory infections. It has been shown that overexpressing mutant forms of MeCP2^T158M^ can ameliorate the Rett phenotype in mice, indicating that the protein is still functional^24^ and that the loss of stability is the major disease driving phenotype of the mutant.

Currently, curative treatment for Rett syndrome is not available due to the complexity of the disease^25^. Recently the first drug specifically developed for Rett syndrome was approved by the FDA^26^. DAYBUE™ reduces brain inflammation by increasing insulin like growth factor 1 (IGF-1). Due to the monogenetic cause of Rett syndrome, gene therapy has been considered as a treatment. Yet, the tight regulation of MeCP2 expression levels and, more importantly, the mosaicism of the disease create a significant challenge for gene therapy approaches.

Here, we established a cell-based phenotypic assay using NIH3T3 cells overexpressing eGFP-tagged MeCP2 or its unstable T158M mutant. The phenotypic difference between both is evident. Wild-type MeCP2 is present in pericentromeric speckles in the nucleus while MeCP2^T158M^ is more diffuse because of impaired DNA binding^24^. We used a high content imaging platform to screen for drugs that are able to correct MeCP2 binding to DNA. In this model system, we screened for small molecules from a repurposing library. More than 3000 compounds were tested, 18 hits were considered positive and recovered at least 25% of the number of speckles in the wild-type cell line with a viability of more than 55%. Among the 18 hits, an histone deacetylase inhibitor (HDACi), PCI-34051, was found to restore MeCP2 speckles to 25% at 10 µM. HDACis have been tested to treat different neurological disorders such as amyotrophic lateral sclerosis (ALS)^27^, Parkinson disease (PD) ^28^ and Rett Syndrome^29,30^. The identification of HDACis in our phenotypic assay validates our screening platform.

## Methods

### Cell culture

All cell lines were cultured in a humidified atmosphere of 5% CO_2_ at 37°C. They were cultured in Dulbecco’s Modified Eagle Medium (DMEM) supplemented with 5% fetal bovine serum (FBS, Gibco) and 0.01% (vol/vol) gentamicin (Gibco) and respective antibiotics for vector selection, 0.005% (vol/vol) blasticidin and 0.001% (vol/vol) puromycin.

### Lentiviral vectors and stable cell lines

shRNA targeting MeCP2 were designed using Snapgene and synthesized at Integrated DNA Technologies (IDT, Belgium). The shRNA was generated by annealing of the sense and antisense oligos. The annealed product was cloned into the pGAE_SFFV plasmid backbone using the Esp3I restriction enzyme. For MeCP2E1-eGFP and MeCP2E1-eGFP^T158M^ back-complementation, plasmids were generated using primers to amplify MeCP2 by PCR and inserted into the phPGK-eGFP-Puro backbone using *XhoI* and *XbaI* restriction enzymes.

NIH3T3 cells stably expressing MeCP2E1-eGFP or MeCP2E1-eGFP^T158M^ were generated by lentiviral vector transduction and subsequent puromycin selection. Stable polyclonal cells were expanded for further selection of monoclonal cell lines. For each cell line, two monoclonal lines were selected and verified by fluorescence microscopy.

### Phenotypic assay

At day 0, 5000 cells from each cell line were plated in a 96-well plate (PhenoPlate 96-well, black, Perkin Elmer) according to the layout in Figure 3A. The cells were incubated for 24 h in 5% CO_2_ and at 37°C. On day 1, 50 µL of compound diluted in cell culture medium was added to each well at a final concentration of 10 µM, compound library provided by CD3 (supplementary table 2). On day 2, the plates were emptied using Bluewasher (Bluecatbio) and the cells were fixed with 4% paraformaldehyde and 5 µg/mL of Hoechst in PBS for 20 min at RT. After fixation PFA was removed and 100 µL of PBS were added. The plates were stored at 4°C. The plates were then imaged using the CX7 (Thermofisher), channel 1-wide-field 386-23: Hoechst 33342, channel 2 – wide-field 488-20: GFP with the 20x objective, with 4 images per well. The images were analyzed using the Cellinsight software (Thermo Fisher). First the software detects the nucleus of each cell in the Hoechst channel. This is used to quantify the number of cells whereas the GFP channel is used to detect and quantify MeCP2 speckles inside the nucleus, previously detected by the Hoechst channel. The small molecules were considered positive if they rescued at least 25% of the number of speckles counted in the wild-type condition.

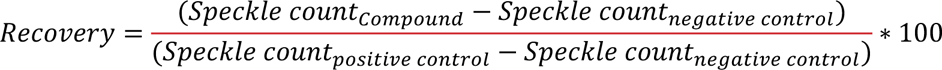

### EC_50_ Determination

To calculate the EC_50_ and CC_50_ of each positive hit, the phenotypic assay was performed with a 3-fold serial dilution of each compound starting at 60 µM and 0.6% of DMSO. The half-maximum effective concentration (EC_50_) was calculated using GraphPad software, 4-way fitting curve, fixing the top at the maximum recovery observed in the wild-type condition. To calculate the half-maximum cytotoxic concentration (CC_50_), we considered the total number of cells per well in the non-treated condition as positive control for viability.

### Protein purification

MiniMeCP2 (aa78-aa309 of MeCP2E2) and miniMeCP2^T158M^ constructs, were produced from pET-MeCP2 constructs in E. *coli* BL21 (see Figure 4). The bacterial culture was grown until an OD of 0.8 (OD = 600 nm)in lysogeny broth (LB) supplemented with 0.5% glycerol, after which expression was induced using 0.25 mM isopropyl β-D-1-thiogalactopyranoside (IPTG) at 18°C for 20 hours. The cells were harvested by centrifugation and lysed using Ni-CAM buffer (25 mM Tris-HCl, 1 M NaCl, 10 µM EDTA, 0.05% β-mercaptoethanol). Then the lysate was sonicated and the supernatant collected after centrifugation for 30 min at 15, 000 rpm. The fusion protein was captured using His-Select Nickel affinity gel [Sigma; P6611] beads and eluted with Ni-CAM buffer supplemented with 250 mM imidazole. Next, the protein was concentrated using Amicon filters [Merck; UFC500324] and run on a Superose 6 10/300 GL size exclusion column.

### Differential Scanning Fluorimetry

The melting temperature of miniMeCP2 and miniMeCP2^T158M^ were measured in the CFX Opus 96 Real-time PCR instrument (Bio-Rad, Belgium) using pre-treated (95^°^C for 10 min to reduce adherence) PCR tubes and optical flat caps (Bio-rad, Belgium). The final reaction volume was 20 µL (10 µM of miniMeCP2 or 20 µM of miniMeCP2^T158M^), 10x of SYPRO™ Orange protein Gel Stain (Thermofisher Scientific) in DSF buffer (50 mM HEPES and 150 mM NaCl at room temperature for 20 minutes. After incubation, the sample was heated progressively from 20°C to 95°C (0.2°C steps). Fluorescence measurements were taken at each step from the SYPRO™ Orange dye to generate a melting curve. The T_m_ of each protein was analyzed using the CFX Maestro software of Bio-Rad.

### Western blot detection

Cells pellets were obtained from 1×10^6^ HEK293T and NIH3T3 cells and dissolved in 100 µL of RIPA buffer, followed by an incubation of 30 min on ice. The supernatant was collected after 10 min of centrifugation at 17000 g at 4 °C. Protein concentration was measured by the bicinchoninic acid (BCA) assay (Thermofisher Scientific). A total of 30 µg of protein extract was loaded on a 4%-12% SDS-PAGE gel and run for 1h at 130 V. The proteins were transferred onto a nitrocellulose membrane (Amersham) using a Turbo blotting protocol from the manufacturer. The membrane was incubated 1 h at room temperature with 4% of skimmed milk in PBS-T (PBS with 0.1% Trinton-X-100). After being washed 3 times with PBS-T the membrane was incubated overnight at 4°C with primary antibodies, 1:500 dilution of rabbit anti-MeCP2 (Cell signalling) and 1:10000 dilution of mouse anti-vinculin (Sigma Aldrich). After the incubation with the primary antibody, the membrane was washed with PBS-T and incubated 1 h at room temperature with the secondary antibodies. The secondary horseradish peroxidase (HRP)-conjugated antibodies used were goat anti-rabbit and goat anti-mouse, both diluted 1:5000 in blocking buffer. The secondary antibody was washed away with PBS-T and stained proteins were detected by chemiluminescence (Clarity western ECL, Bio-rad). The western blot signal was acquired using Amersham Image Quant8000.

## Results

### Construction of monoclonal cell lines overexpressing MeCP2E1-eGFP or MeCP2E1-eGFP^T158M^

The differential phenotype of MeCP2 and MeCP2^T158M^ has been studied in murine NIH3T3 ^12,13^. Wild-type MeCP2 forms clear pericentromeric speckles while MeCP2^T158M^ displays less speckles due to impaired affinity to chromatin^12^. Speckles are known to represent liquid phase condensates ^12^. To identify compounds targeting human MeCP2 we initially aimed to detect if in human cell lines such as HEK293T a different phenotype between wild-type and mutant MeCP2 could be visualized. Unfortunately, no speckled pattern was present in HEK293T and the difference between the wild-type and MeCP2^T158M^ was not easily spotted in this cell line. Instead, we depleted endogenous mouse MeCP2 by shRNA knock down and overexpressed the human isoform C-terminally tagged with eGFP in murine NIH3T3 cells (Figure 2.) We generated isoform specific cell lines to compare MeCP2E1 with MeCP2E2. In Figure 2 human MeCP2E1-EGFP adopts the typical speckled pattern of endogenous mouse MeCP2. Moreover, the difference between MeCP2 and MeCP2^T158M^ speckles was clear for both isoforms, indicating that the eGFP-tag does not affect the binding of MeCP2 to DNA. This difference encouraged us to establish a phenotypic assay.

**Figure 1.**
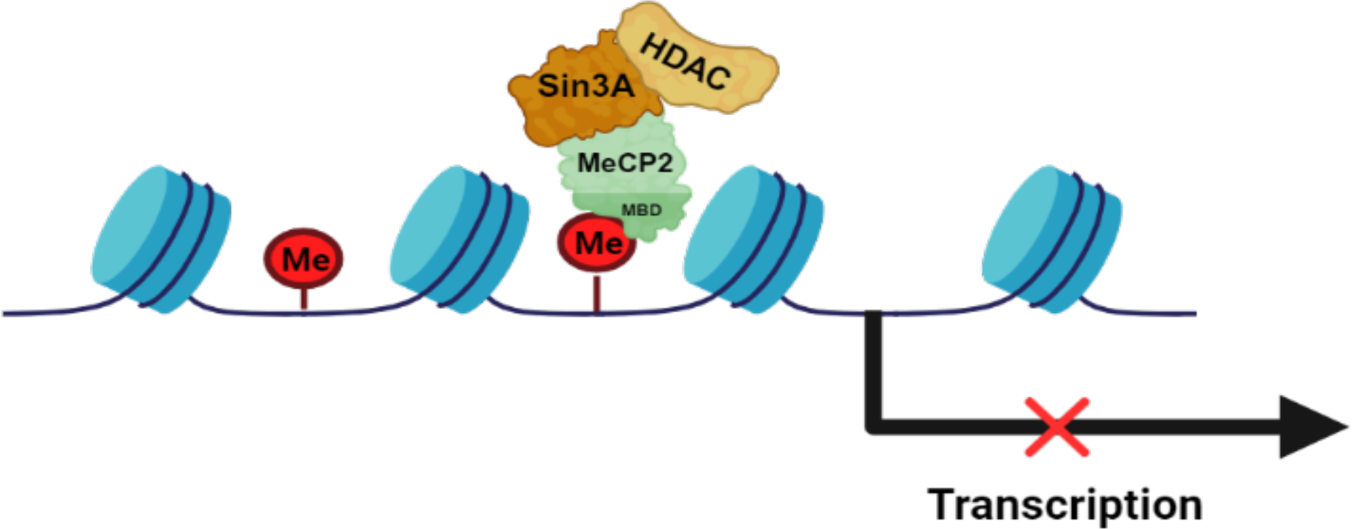
MeCP2 binds to methylated DNA (Me) via MBD. Sin3A and HDAC form a triple complex with MeCP2 blocking transcription.

**Figure 2.**
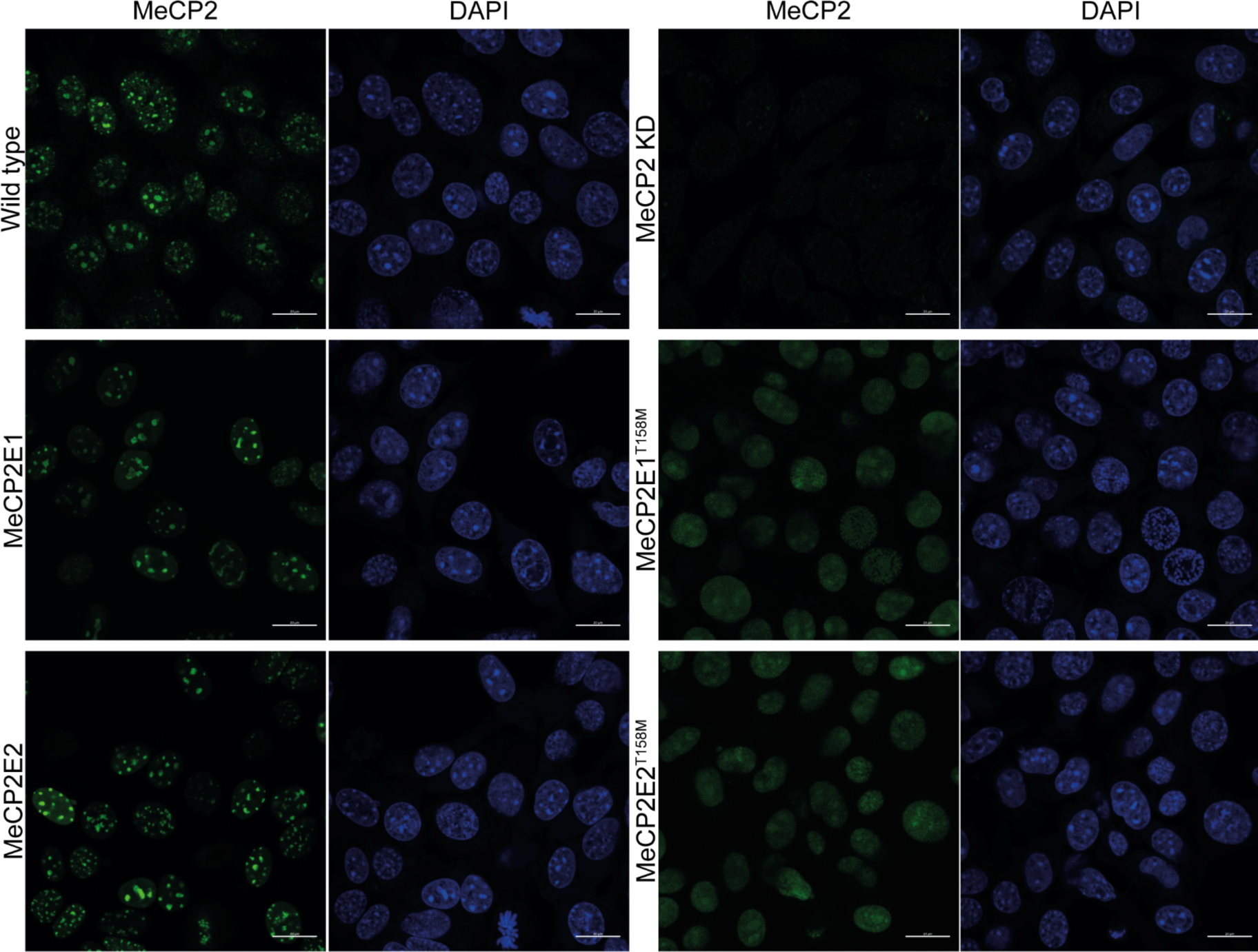
MeCP2 phenotype in various cell lines. Confocal microscopy images show cellular distribution of MeCP2 in NIH3T3 wild–type and MeCP2 knock down (KD) cells. NIH3T3 cells overexpressing MeCP2E1-GFP, MeCP2E1-T158M-eGFP, MeCP2E2-eGFP or MeCP2E2-T158M-eGFP display different distributions of MeCP2. The cell lines overexpressing MeCP2-eGFP are monoclonal cell lines. Endogenous MeCP2 was detected with rabbit anti-MeCP2 antibody (1:500) and anti-rabbit alexa 488 secondary antibody (1:1000) (upper row) while eGFP-tagged MeCP2 was detected in the 488 nm channel in green (first column) and DAPI in the 405 nm channel (blue). Scale bar – 20 µm.

The phenotypic assay accurately assesses the speckled pattern of wild-type MeCP2E1 and MeCP2E1^T158M^. MeCP2E1 forms distinct and clear pericentromeric speckles in the nucleus, whereas MeCP2E1^T158M^ is distributed dispersedly (Figure 2). Stable NIH3T3 cell lines expressing MeCP2E1-eGFP or MeCP2E1-eGFP^T158M^ were generated using lentiviral vector transduction and puromycin-based selection, followed by clone selection to avoid heterogeneous expression levels. Confirmation of the phenotype was achieved through fluorescence microscopy and quantification of speckles per nucleus, which revealed an average of 13.6 ± 0.6 speckles per cell for MeCP2E1-eGFP and only 1 ± 0.5 speckle for MeCP2E1-eGFP^T158M^ (Figure 3D).

**Figure 3.**
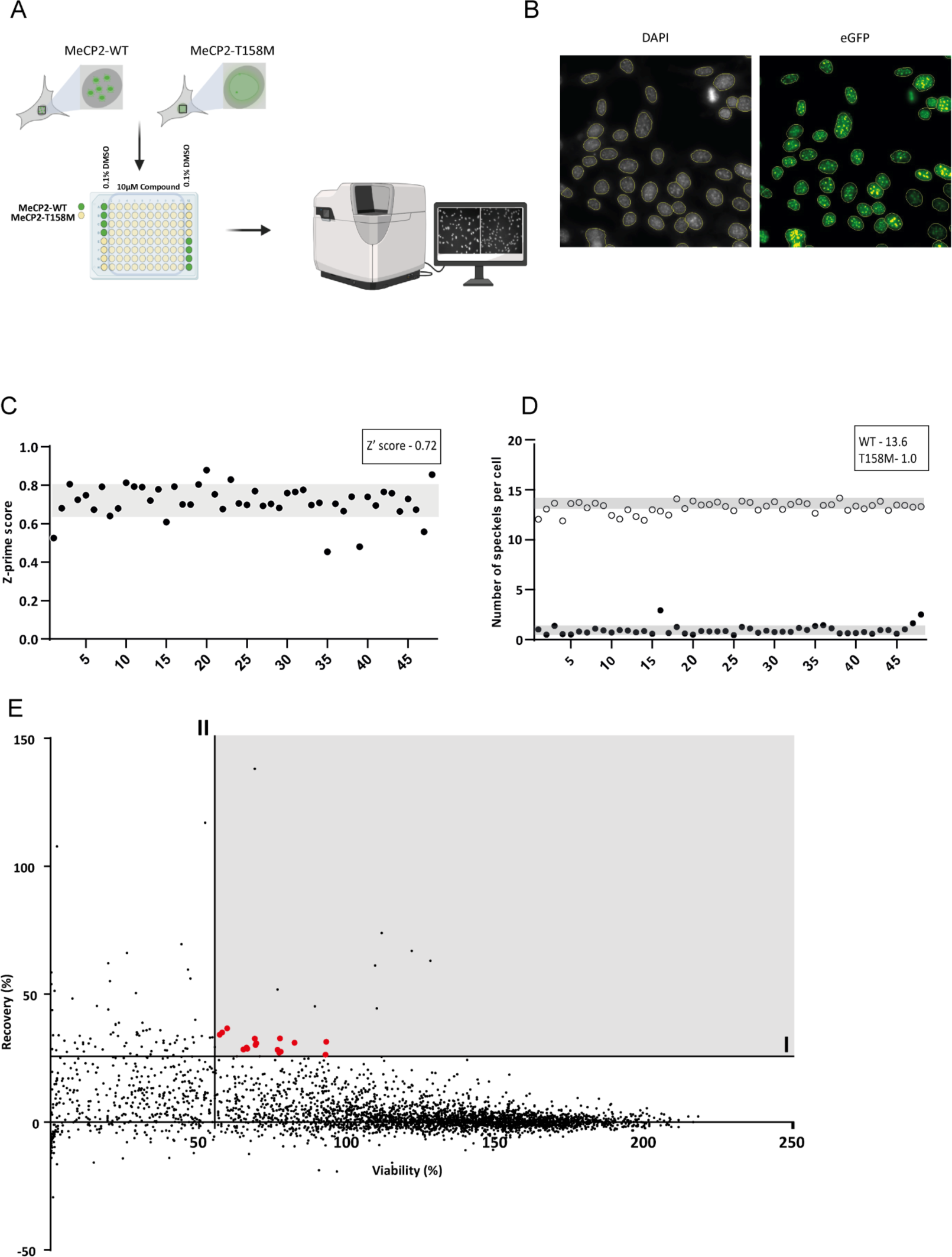
High content screen. **A)** Schematic representation of the high content screen pipeline. The cell lines NIH3T3-MeCP2E1-eGFP and NIH3T3-MeCP2E1-eGFP^T158M^ were seeded in 96-well plates, the wells with wild-type MeCP2 are displayed in green, those with MeCP2^T158M^ in yellow **B)** Representative picture of DAPI staining and delimitation of nucleus (yellow line). Representation of eGFP detection in the nucleus and respective spot detection (yellow dots). **C)** Individual representation of Z’-score of each plate. The average Z’-score of the screen was 0.72 ± 0.08 **D)** Number of speckles detected in the wild-type control (positive control) and the T158M cell line (negative control) per plate**. E)** Representation of the individual values for each compound for Viability (%) X axis and recovery (%). Horizontal line (I) – Compounds that recover more than 25% of the wild-type phenotype. Vertical line (II) – Compounds that show 25% recovery and at least 55% viability. The grey area represents the compounds considered as positive hits. Red dots represent <u>the18 compounds considered as positive hits.</u>

As a positive control in the screen, NIH3T3-MeCP2E1-eGFP cells were seeded in wells A1 to D1 and E12 to H12, while the negative control NIH3T3-MeCP2E1-eGFP^T158M^ cells were seeded in wells E1 to H1 and A12 to D12. Furthermore, all wells were treated with DMSO to a final concentration of 0.1% to account for the solvent the compounds were diluted in. The remaining wells of each plate were filled with NIH3T3-MeCP2E1-eGFP^T158M^, and the small molecules were added at a final concentration of 10 µM.

We screened 3572 small molecules from a Drug-Repurposing Library of the Centre for Drug Design and Discovery (CD3, Leuven) to assess their ability to correct MeCP2^T158M^ binding to DNA and facilitate the formation of pericentromeric domains. To ensure reliable results, controls were placed on the outer wells of the plate, and the Z’-factor was calculated using mean values and standard deviation of speckles per cell in NIH3T3-MeCP2E1-eGFP and NIH3T3-MeCP2E1-eGFP^T158M^ (Figure 3C). The assay demonstrated a robustness with an average Z’-prime score of 0.7 ± 0.1. Small molecules were considered positive hits only if they showed at least 25% recovery of the wild-type speckle count with over 55% cellular viability. Out of the 3572 small molecules, 118 showed at least 25% rescue of the phenotype, but only 28 passed toxicity criteria. To ensure validity, individual inspection of images led to the removal of 10 compounds from the selection due to saturation in the GFP channel. In the final list of 18 compounds, the range of rescue varied from 25.7% up to 36.6%, with cell viability ranging from 55.5% to 99.2% (Table 1).

**Table 1.**
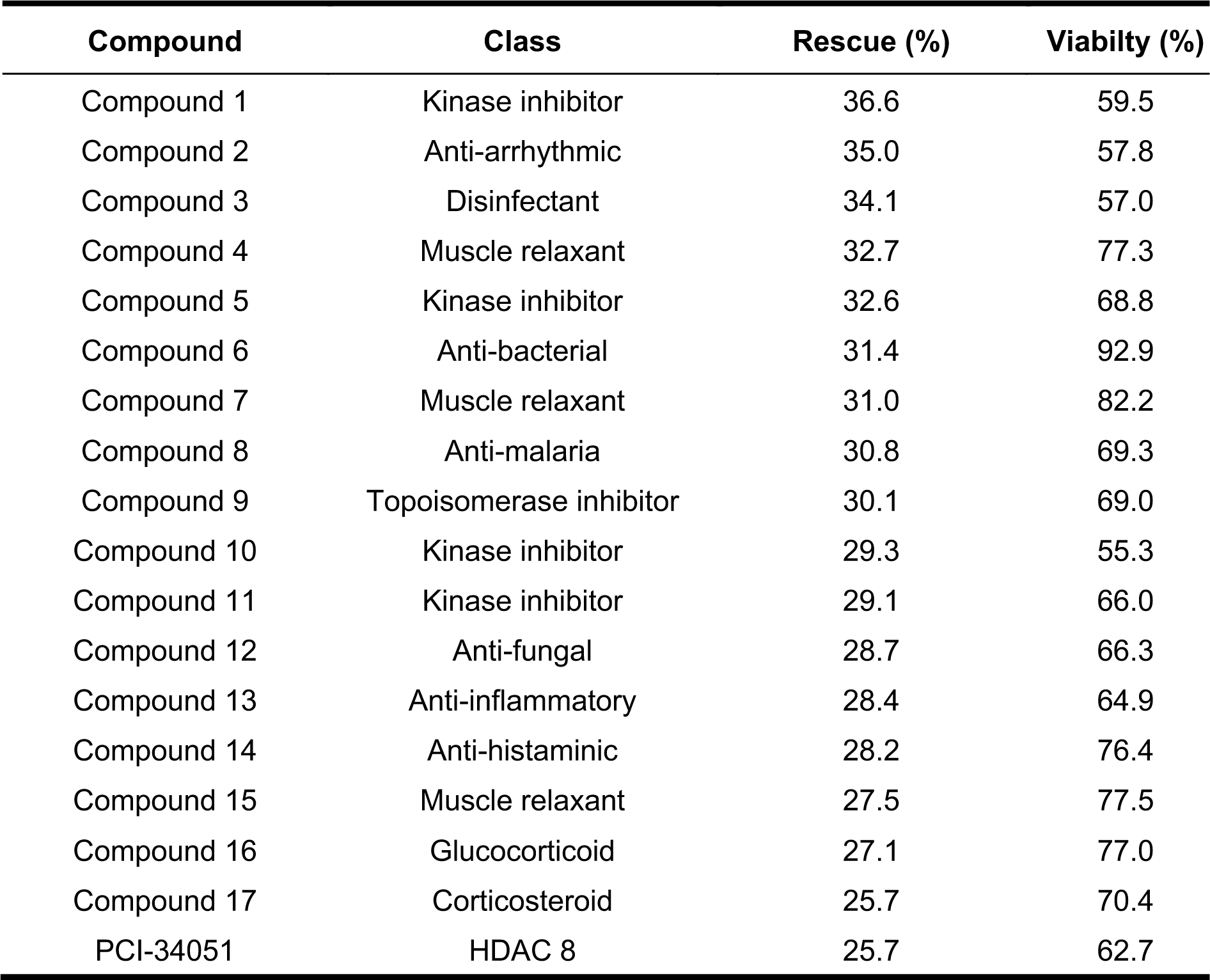
List of positive hits in high content screening. Rescue represents the number of speckles in % relative to wild-type. Viability in % relative to a DMSO control.

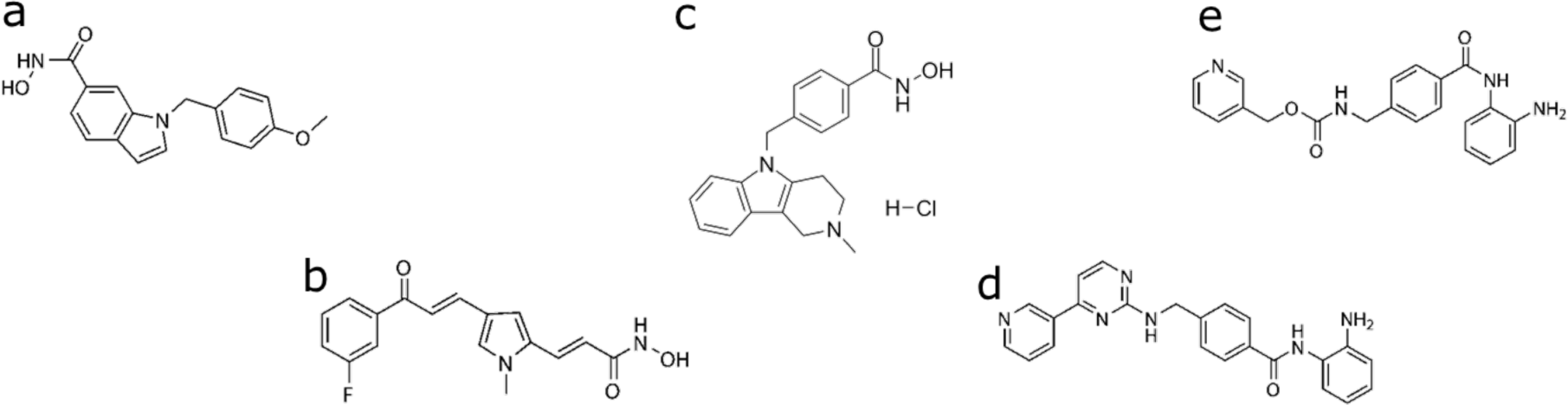

### Compounds demonstrate a dose-response activity rescuing MeCP2 expression

To validate the positive hits from the high content screen, we determined their dose-response using a ten step 3-fold serial dilution of each compound, starting at 60 µM. In parallel, we included four HDACis detailed in Table 2, which were initially classified as negative in the high content screen. The objective was to comprehend whether HDACis have any impact on MeCP2’s DNA binding affinity, despite their lack of positive effects in the initial screening. In the first screening, several compounds demonstrated activity. However, TBA did not reach the required 25% threshold, and additionally demonstrated toxicity. Mocetinostat displayed activity but was associated with toxicity. Entinostat showed a recovery below the 25% threshold, yet it did not exhibit any toxicity allowing us to increase the concentration during dose response. Despite MC-1568 meeting both thresholds, it was initially excluded due to its autofluorescence. However, it was included in the dose response to explore the possibility of reduced autofluorescence at lower concentrations without compromising efficacy. This decision turned out to be correct as we observed an increase in the number of speckles at very low concentration, indicating a positive effect at reduced autofluorescence levels.

**Table 2:**
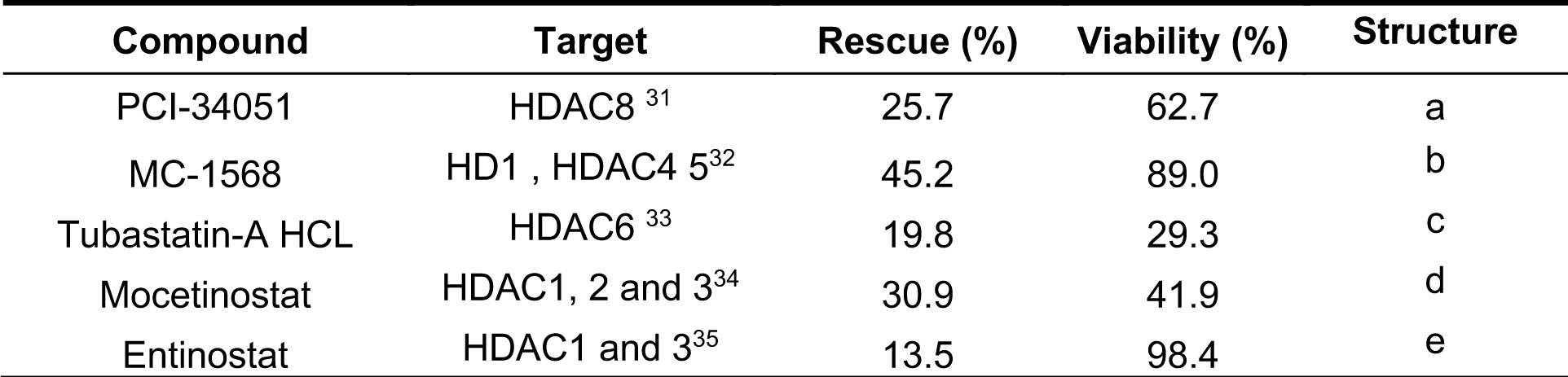
List of HDACis included in the dose response (DR) assay and corresponding structures.

Considering that MeCP2 has two isoforms with different expression patterns and functions^8^, we also included both MeCP2E1 and E2 in the dose response assay to evaluate if the compounds have similar effects on both isoforms.

As described in Table 3, for most compounds a dose response (EC_50_) could be determined. Despite showing a dose response, the maximum recovery observed was 6 speckles per cell, corresponding to less than 50% of the wild-type, thus we calculated the EC_50_ taking that number as maximum recovery possible. The EC_50_ of the positive hits for MeCP2E1^T158M^ ranged from 0.6 to 113.78 µM. For the specific class of the HDACis there was less variability with a range from 6.09 µM to 14.67 µM. The IC_50_ of entinostat on the contrary was only 113.78 µM but it shows high variability (Table 3). For MeCP2E2^T158M^ similar results were obtained, the EC_50_ for each compound was in the same range as with MeCP2E1^T158M^ (Supplementary Table 1). The toxicity levels was consistent across all cell lines tested, however the selectivity index is low, indicating that activity is associated with toxicity.

**Table 3.** EC_50_ and CC_50_ values (µM) in dose response assay for NIH3T3-MeCP2E1 and NIH3T3-MeCP2E1^T158M^. NIH3T3-MeCP2E1 n = 2, NIH3T3-MeCP2E1^T158M^ n= 3. Compounds 6,11,16 were not available for retesting (N/A).

|  | E1<sup>T158M</sup> |  |  |  | E1 |  |  |
| --- | --- | --- | --- | --- | --- | --- | --- |
|  | EC<sub>50</sub> |  | CC<sub>50</sub> |  | Selectivity index | CC<sub>50</sub> |  |
|  | Average | StDev | Average | StDev |  | Average | StDev |
| Compound 1 | 1.80 | 1.63 | 1.84 | 0.91 | 1.02 | 2.59 | 2.14 |
| Compound 2 | 8.22 | 7.55 | 8.66 | 5.55 | 1.05 | 9.81 | 6.46 |
| Compound 3 | 13.23 | 5.76 | 14.85 | 6.55 | 1.12 | 9.02 | 3.47 |
| Compound 4 | 5.57 | 2.13 | 6.84 | 3.07 | 1.23 | 5.80 | 4.21 |
| Compound 5 | 3.46 | 1.29 | 0.83 | 0.37 | 0.24 | 0.61 | 0.34 |
| Compound 6 | N/A | N/A | N/A | N/A | N/A | N/A | N/A |
| Compound 7 | 15.78 | 11.87 | 12.14 | 0.75 | 0.77 | 12.44 | 8.83 |
| Compound 8 | 51.13 | 38.27 | 36.82 | 12.46 | 0.72 | 29.31 | 0.00 |
| Compound 9 | 0.60 | 0.81 | 2.34 | 2.06 | 3.92 | 4.00 | 2.18 |
| Compound 10 | 4.47 | 3.78 | 4.39 | 2.39 | 0.98 | 5.49 | 4.40 |
| Compound 11 | N/A | N/A | N/A | N/A | N/A | N/A | N/A |
| Compound 12 | 6.41 | 4.31 | 6.93 | 0.97 | 1.08 | 6.52 | 4.75 |
| Compound 13 | 6.79 | 1.64 | 7.52 | 1.86 | 1.11 | 5.74 | 2.69 |
| Compound 14 | 13.67 | 5.19 | 8.73 | 0.72 | 0.64 | 8.56 | 6.04 |
| Compound 15 | 2.85 | 1.76 | 2.18 | 1.01 | 0.76 | 3.48 | 2.85 |
| Compound 16 | N/A | N/A | N/A | N/A | N/A | N/A | N/A |
| Compound 17 | 15.50 | 13.35 | 5.85 | 3.72 | 0.38 | 28.20 | 0.00 |
| Mocetinostat | 8.47 | 0.55 | 3.89 | 0.33 | 0.46 | 3.59 | 1.99 |
| Entinostat | 113.78 | 92.66 | 35.29 | 7.66 | 0.31 | 16.94 | 10.38 |
| PCI-34051 | 14.67 | 10.15 | 16.97 | 5.36 | 1.16 | 11.71 | 2.94 |
| Tubastatin-A | 7.45 | 4.64 | 2.53 | 0.52 | 0.34 | 3.44 | 0.97 |
| MC-1568 | 6.09 | 1.28 | 11.20 | 1.71 | 1.84 | 13.54 | 6.41 |

The hits from the phenotypic screen were subjected to a differential scanning fluorimetry (DSF) assay to investigate their impact on the melting curve of miniMeCP2^T158M^. We purified a truncated version of MeCP2, named mini-MeCP2, which includes MBD, ID and TRD (Figure 4A). Notably, the wild-type mini-MeCP2 exhibited a melting temperature (Tm) of 45°C, while the mutant mini-MeCP2^T158M^ displayed a significantly reduced Tm of 32.5°C (Figure 4B), indicating decreased protein stability for the mutant MeCP2. Using this assay, we assessed the effect of the hits from the phenotypic screening. Among the 15 compounds tested, only compound 13 exhibited a significant increase in the melting temperature, indicating its direct binding to mini-MeCP2^T158M^. In contrast, the HDAC inhibitors that target class I HDAC, Entinostat and Mocetinostat showed a significant decrease in the melting temperature of miniMeCP2^T158M^ when tested against wild-type mini-MeCP2. Neither compound 13 nor HDAC inhibitors displayed a significant effect on the wild-type protein, indicating that compound 13 specifically targets the mutant protein.

**Figure 4.**
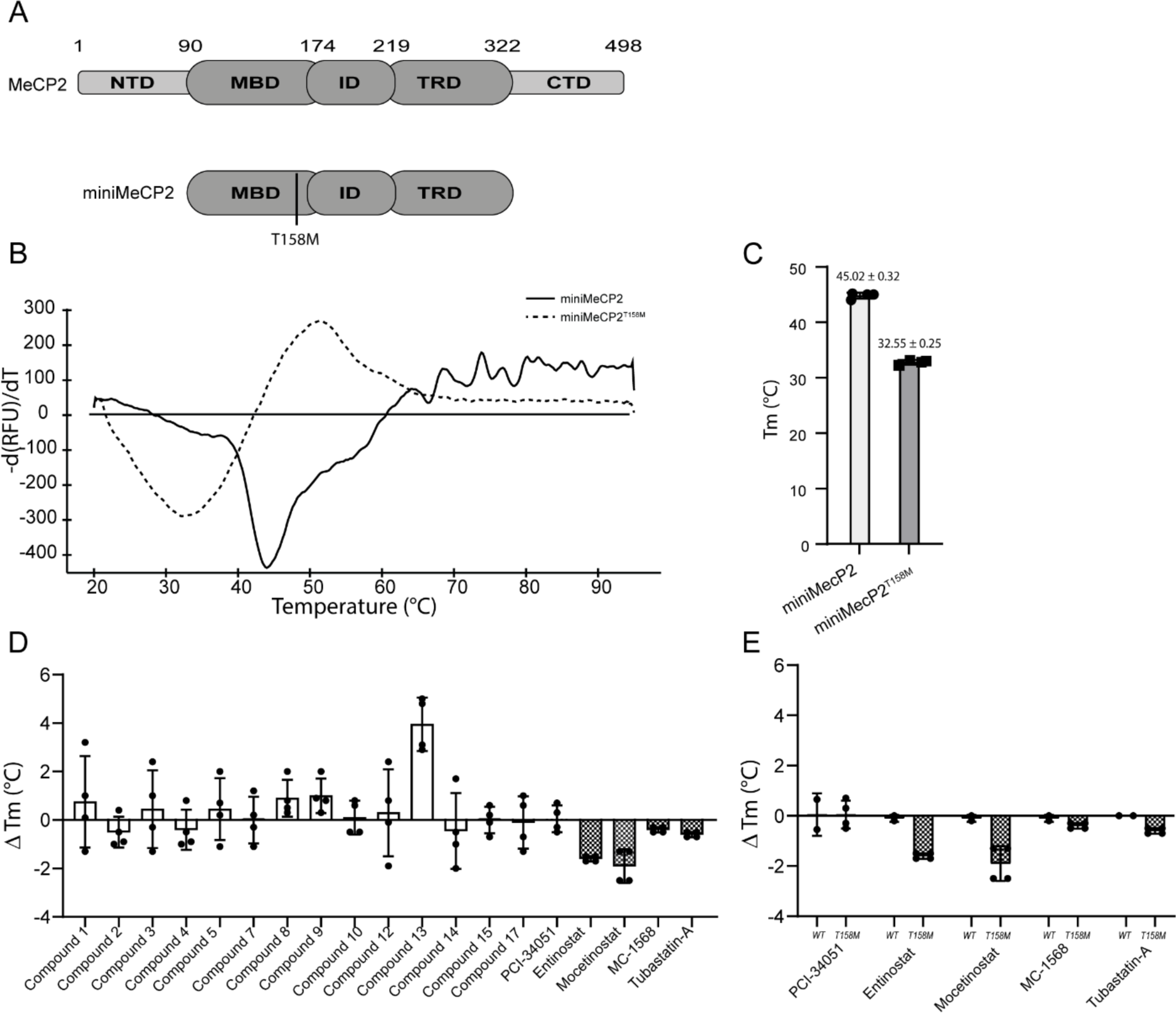
Protein stability of miniMeCP2^T158M^ is affected by Compound 13.1. **A)** Schematic representation of MeCP2 full length and miniMeCP2. **B)** Melting curve of miniMeCP2 and MiniMeCP^T158M^ in DSF assay, negative peak of fluorescence represented. **C)** Melting temperature (Tm °C) of miniMeCP2^wt^ and miniMeCP2^T158M^, calculated from the melting curve in the SYPRO assay. **D)** Graphical representation of thermal shift (°C) of miniMeCP2^T158M^ when incubated with 10 µM of compound relative to DMSO control. **E)** Effect of 10 µM of five different HDACis in WT and T158M mini MeCP2.

### HDACis Affect MeCP2 Expression

Considering that the HDACis did not increase the melting temperature of MeCP2^T158M^, we anticipated that they might influence its expression levels. To investigate this, we treated HEK293T and NIH3T3 wild-type cells with 10 µM of each HDACi. By comparing both cell lines, one with mouse MeCP2 and the other with human MeCP2, we aimed to test the impact of HDACis in cells expressing both isoforms of the respective MeCP2 variants. Since different HDACis target various HDAC classes, this allowed us to explore the effects of individual HDACs on the expression of MeCP2. As shown in Figure 5, treatment with HDACis resulted in reduced MeCP2 levels, as reported before^30^. Treatment with Tubastin-A, Mocetinostat or Entinostat reduces MeCP2 levels in both cell lines. In contrast MC-1568 shows a reduction of MeCP2 levels in NIH3T3 cell line but no effect in HEK293T cells, whereas PCI-34051 shows an effect in HEK293T but not in NIH3T3, indicating distinct mechanisms in these cell lines.

**Figure 5.**
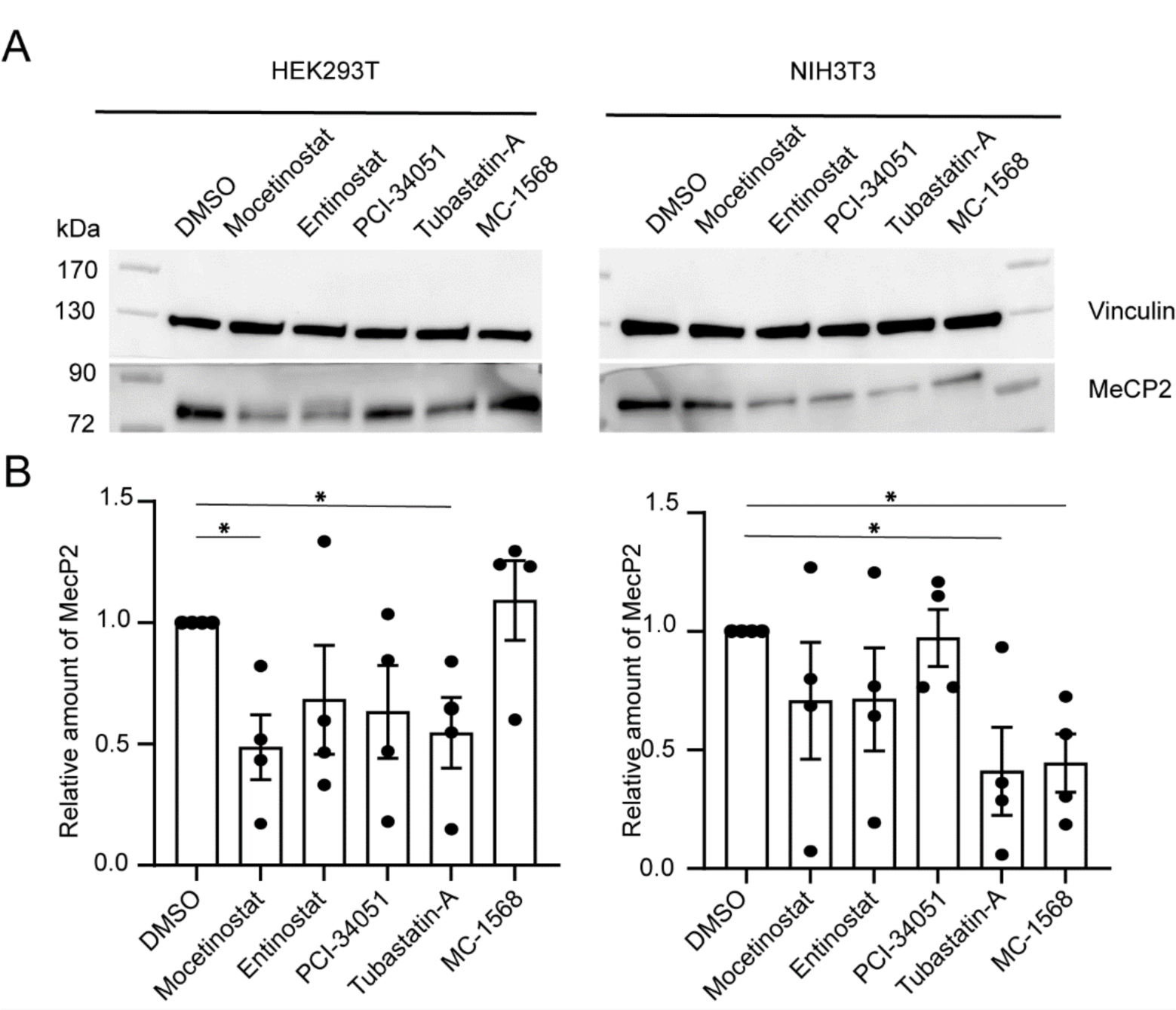
HDAC inhibitors affect MeCP2 protein levels. **A)** Representative blots of the effect of 10 µM of HDACi during 24 h on MeCP2 levels in HEK293T (left) or NIH3T3 (right). **B)** Densitometry measurements of MeCP2 levels, normalized to the DMSO control. Data represented as mean ± SEM of 4 experiments (each individual dot). Mann-Whitney test was used to assess statistical differences. * *p* value < 0.05.

## Discussion

Finding a specific treatment for Rett Syndrome is a high unmet medical need and has been the aim of many research groups in both academia and industry in recent years. Despite the significant effort and advances, there is still no curative therapy focused on correcting the impact of RTT mutations on the function of MeCP2. Recently, FDA approved the first specific treatment for RTT patients, trofinetide, which is an analogue of insulin like growth factor 1 (IGF-1), acting as an anti-inflammatory agent.^26^ Although it is considered a specific RTT treatment, trofinetide does not target MeCP2, the cause of RTT. Other clinical trials are ongoing trying to deliver a functional version of MeCP2 using adeno-associated viral vectors (AAV)^36^. The gene therapy approach seems to face many obstacles due to the mosaicism of RTT, specifically targeting the unhealthy cells and delivering the correct dose to avoid MeCP2 duplication syndrome (MDS)^37^. MDS is an extremely rare condition that primarily affects boys, as they only have one X-chromosome. Girls can also be affected but often exhibit milder symptoms due the mosaic effect of the X-chromosome. RTT is caused by more than 800 different mutations in MeCP2, leading to pleiotropic effects, such as malfunction, reduced expression or expression of non-functional truncated MeCP2 proteins. Due to these pleiotropic effects, finding a single drug to cure RTT will be a difficult if not impossible endeavor. Here, we focused on the most common single point mutation, T158M, accounting for almost 10% of all RTT patients^18^.

Lamonica *et al* ^24^ showed that overexpression of MeCP2^T158M^ in mice rescues the RTT phenotype, indicating that the protein retains its affinity for DNA, yet cannot exert this function sufficiently due to reduced stability, which has also been suggested by other authors ^20,38,39^. We established a phenotypic screen to identify small molecules that can revert this phenotype and allow MeCP2^T158M^ to form condensates similar to wild-type. The crucial part of this work was to create a cell model in which the condensate forming capability of MeCP2 and MeCP2^T158M^ could be discriminated. Initially, we tried to establish the assay in human cell lines however, the difference observed between MeCP2 and MeCP2^T158M^ in human cell lines was less clear than in mouse cell lines since the speckled pattern is mouse specific as described earlier^13^. The speckled pattern displayed by mouse cells is due to heterochromatin arrangement independently from MeCP2^13^. Despite the high identity of human and mouse MeCP2 we engineered NIH3T3 (mouse fibroblasts) to express human MeCP2 to ensure that the identified hits indeed act on the human protein.

In the phenotypic assay, the number of MeCP2 speckles in the nucleus of NIH3T3 was quantified. MeCP2^T158M^ was classified as having abolished clustering, meaning that it still binds to DNA but without the clear speckled pattern^12,21^. To reduce the individual experimental steps in the screening process, and thus minimize variability and duration of the assay, we opted to tag human MeCP2 with eGFP. We explored both N-terminal and C-terminal eGFP tags and there was no difference in the speckle formation. However, since MeCP2E1 and MeCP2E2 differ only in the N-terminal domain that determines protein stability by the N-end rule, we decided to continue with C-terminal tag to avoid masking the difference between the two isoforms. To identify compounds rescuing MeCP2 function, a repurposing library with compounds from different chemical classes was screened. From the 3572 compounds from, 18 hits were identified. To consider a compound as a hit, a rescue of at least 25% of the number of speckles observed in the wild-type was required. Nonetheless, around 0.5% of compounds were positive, in line with other similar screening assays^40^. Those 18 hits that rescued at least 25% of the phenotype provided strong proof of concept that our screening strategy allows to identify hits modifying the phenotype observed in NIH3T3 cells.

Interestingly, the compounds considered positive belong to very different classes, including kinase inhibitors, muscle relaxants and HDAC inhibitors (Table 1), reflecting the diverse nature of our repurposing library and suggesting that correcting MeCP2^T158M^ abolished clustering can be achieved via multiple pathways including direct binding to MeCP2^T158M^.^21^

In our assay, we compared both isoforms carrying the T158M mutation that affects DNA affinity and disrupts the speckle formation. When we tested the positive hits and the additional HDACis in the dose response assay we could show that all compounds display a dose response curve demonstrating the specificity of the compounds. The EC_50_ value was in general lower for MeCP2E2^T158M^, which can be explained by the higher affinity of MeCP2E2 to DNA that can be increased with chaperone-like molecules or by affecting the post-translational modification of MeCP2^5^. ^41^

Our main goal was to find chaperone-like small molecules that stabilize MeCP2^T158M^, hence we investigated whether the compounds affect the folding of the protein. We decided to use mini-MeCP2, that only contains MBD, ID and TRD, avoiding the differences between isoforms. The wild-type mini-MeCP2 showed a higher melting temperature than MeCP2^T158M^, as expected. The significant difference of 13°C, creates a window to measure compounds that affect the melting temperature of mini-MeCP2^T158M^. Interestingly, compound 13 increased the melting temperature of miniMeCP2^T158M^ with 5°C implying a direct binding at 10 µM.

By screening the repurposing library, HDAC inhibitors were identified as one promising class of compounds. This is in line with previous reports, Trichostatin and Tubastatin-A, have been tested in MeCP2 knockout mice showing improvement in the phenotype. While HDAC inhibitors are known to regulate gene expression, we observed that compound 13, Mocetinostat and Entinostat affected the melting temperature of miniMeCP2^T158M^ suggesting that they can have both direct and indirect effects on MeCP2. Previous studies with the generic HDAC inhibitor Trichostatin, have shown that HDACis hyperacetylate histones in non-neuronal cells and decrease the levels of MeCP2^30^. Although this seems paradoxical to what we are pursuing, the chromatin binding affinity is significantly increased due to a decrease in the phosphorylation of MeCP2-S164^30^, which can explain the increase in the number of speckles detected in MeCP2^T158M^ cell lines. Our results show that TBA reduces MeCP2 levels as described for Trichostatin. However, it is worth noting that while Trichostatin is a generic HDACi, TBA specifically targets HDAC 6, a class IIb HDAC. TBA has been tested in the context of RTT. Treatment of MeCP2^308/y^ mice resulted in significant improvement in the mouse behaviour and survival. Although TBA targets HDAC6, the function of HDAC6 still needs further investigation. Recent studies associate HDAC6 with α-tubulin, IFNαR and hepatic metabolism^41^. TBA increases α-tubulin acetylation, promoting axonal trafficking by increasing microtubule stability^18^. In MeCP2 knockout neurons BDNF trafficking is deficient but can be restored with TBA ^18^. The effect of Tubastatin in MeCP2^308/y^ mice suggests that HDACis could be used to treat RTT. Our results indicate moreover that the mechanism of action of this specific compound is very similar in both human and mouse cell lines.

The other HDACis tested target Class I HDACs (HDAC1, 2 and 3) and HDAC8 that is evolutionary closely related ^42,43^. The HDAC1, 2 and 3 inhibitors Mocetinostat and Entinostat showed similar effects in both cell lines. HDAC1 and 2 form a complex with Sin3A^44^, while HDAC3 is recruited by the N-CoR/SMRT corepressor complex that also binds MeCP2. MeCP2 has been described to interact indirectly with HDACs using Sin3A as a bridging element. The complex formed increases the deacetylation of the chromatin, repressing gene transcription^45^. As mentioned previously HDACis can affect MeCP2 levels. It is possible that upon loss of binding partners MeCP2 becomes less stable, increasing the degradation process.

On the other hand, HDAC8 functions without known binding partners, the results from PCI-34051 are more complex to interpret. However, PCI-34051 has an IC_50_ of 10 nM, 1000 fold lower than the concentration used in our experiment, at such high concentration, it can also target other HDACs, and PCI-34051 may also affect HDAC1 and HDAC6 at least in HEK293T.

HDACis can improve the RTT phenotype as shown before^29,30^. However, in our assay the selectivity index is very narrow, which is consistent with the pleiotropic effects of HDACi, thus treating RTT with HDAC inhibitors maybe not be realistic since HDACs are involved in many different pathways. Currently HDACi are used to treat different cancers such as lymphoma or myeloma or neurological disorders such as epilepsy or bipolar disorders^46^.

In conclusion, we established and validated a screening platform to identify compounds that rescue MeCP2^T158M^ speckles. This platform is amenable to screening large chemical libraries. Apart from drug discovery for RTT, hits may help to understand how different cellular pathways affect MeCP2 and MeCP2 mutants.

## Data Availability

The datasets generated during the current study are not publicly available due to intellectual property protection but are available from the corresponding author on reasonable request.

## Authors contribution

Conception and design, Zeger Debyser and Frauke Christ. Material preparation and data collection Rodrigo Lata, Marnik Nijs, Liesbeth Steegmans and Rannie Kellens. Data analysis Rodrigo Lata, Marnik Nijs, Hugo Klaassen and Matthias Versele. Writing original draft, Rodrigo Lata, Zeger Debyser. All authors read and approved the final manuscript.

## Acknowledgements

I acknowledge Siska Van Belle, Paulien van de Velde, Saskia Lesire and Yannick Hoogvliets for excellent technical assistance.

---

**Figure S1:**
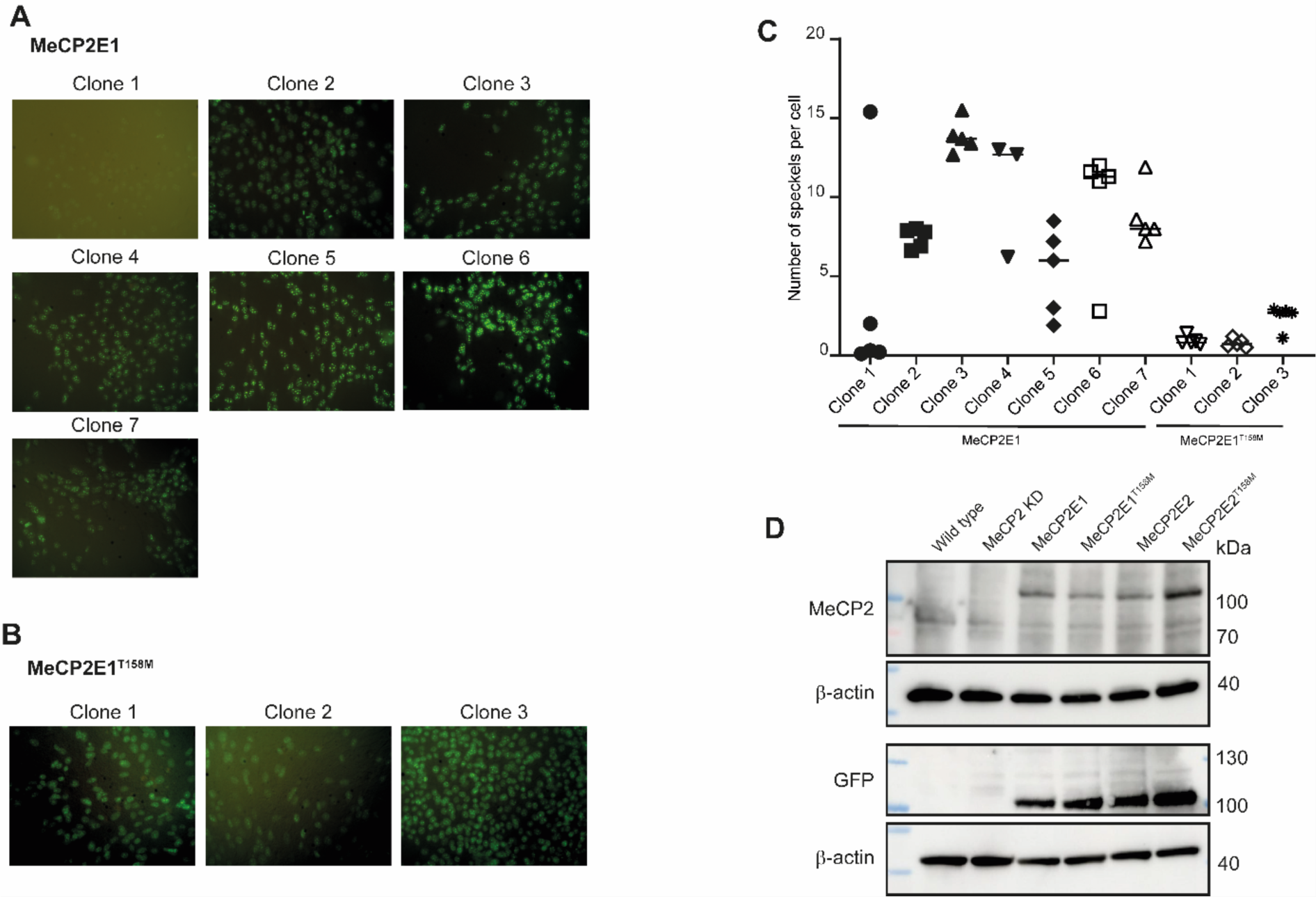
NIH3T3 MeCP2-E1 and MeCP2E1T158M screening for monoclonal lines. Polyclonal lines were seeded in 96-well plates at a density of 1 cell per well. After two weeks each well was checked to confirm the growth of single clones. The wells with one single clone were expanded for further analyses. **A)** Visualization of MeCP2-eGFP localization in NIH3T3MeCP2E1 monoclonal cell lines. **B)** Visualization of MeCP2T158M-eGFP localization in NIH3T3MeCP2E1T158M monoclonal cell lines. These images were captured using a Zeiss fluorescent microscope equipped with a 40x objective lens. **D)** Quantification of number of speckles per cell in NIH3T3-MeCP2E1-eGFP and NIH3T3-MeCP2E1T158M-eGFP. **E)** Detection of endogenous MeCP2 and overexpressed MeCP2-eGFP tagged in NIH3T3 with anti-MeCP2 antibody and anti-GFP antibody. b-actin used as loading control.

**Figure S2:**
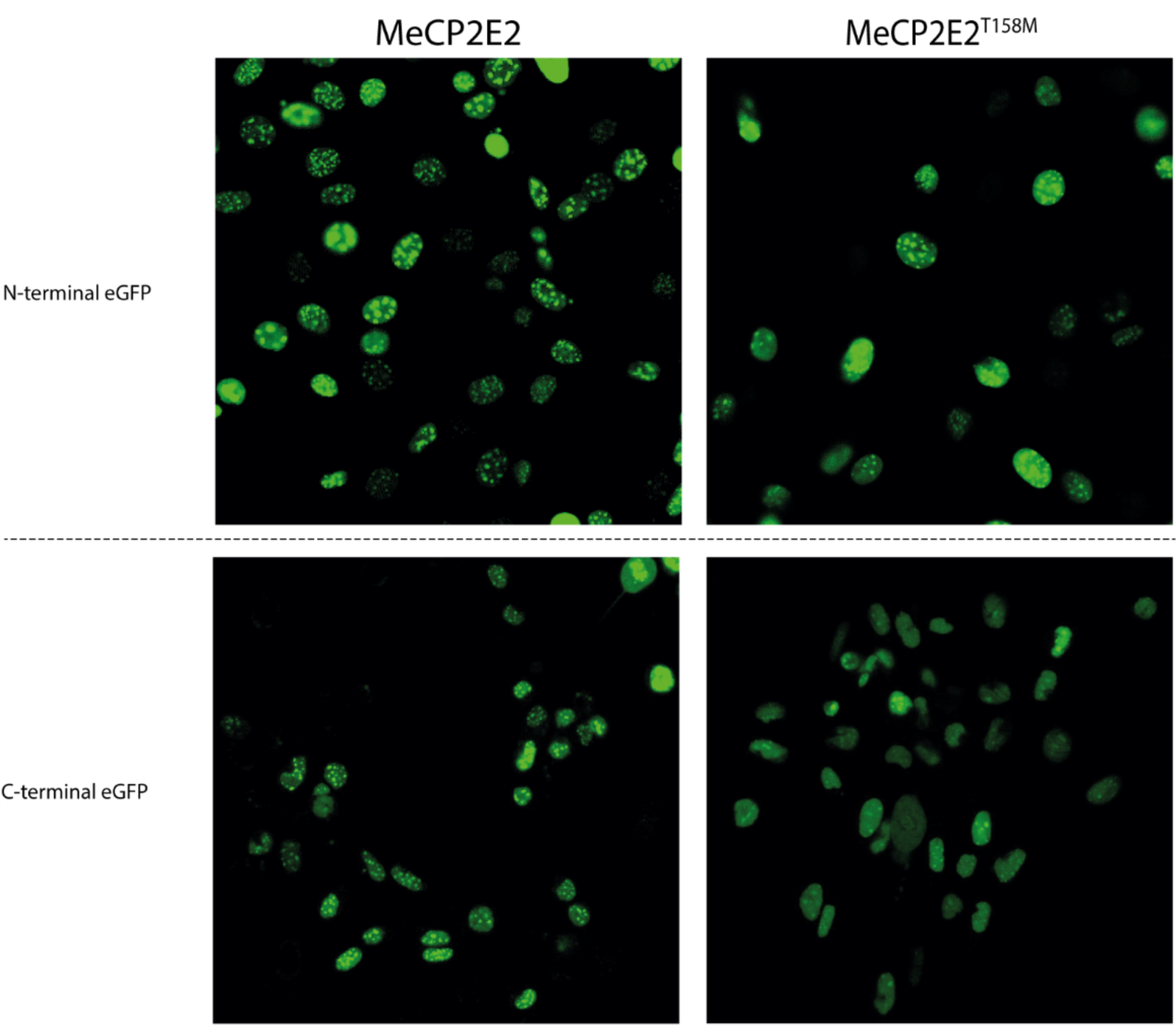
Comparison between N-Terminal and C-Terminal eGFP tag. Representative picture of polyclonal cell lines, NIH3T3E2 and NIH3T3E2T158M with N-Terminal or C-Terminal eGFP tag, images acquired with confocal microscope Zeiss LSM880 with 40x oil objective.

**Table S1.** EC_50_ and CC_50_ values from dose response assays for NIH3T3-MeCP2E2 and NIH3T3-MeCP2E2^T158M^. NIH3T3-MeCP2E1 n = 1, NIH3T3-MeCP2E1^T158M^ n= 1. Compounds 6,11,16 were not available.

|  | E2<sup>T158M</sup> |  |  |  |  | E2 |  |
| --- | --- | --- | --- | --- | --- | --- | --- |
|  | EC<sub>50</sub> |  | CC<sub>50</sub> |  | Selectivity index | CC<sub>50</sub> |  |
|  | Average | StDev | Average | StDev |  | Average | StDev |
| *Compound 1* | 5.03 | 2.92 | 8.15 | 1.66 | 1.62 | 4.85 | 1.89 |
| *Compound 2* | 8.86 | 7.41 | 20.19 | 7.49 | 2.28 | 14.97 | 11.55 |
| *Compound 3* | 19.56 | 0.23 | 12.18 | 6.16 | 0.62 | 15.52 | 3.19 |
| *Compound 4* | 4.57 | 2.23 | 8.39 | 0.31 | 1.84 | 8.33 | 1.33 |
| *Compound 5* | 33.65 | 31.85 | 1.92 | 0.80 | 0.06 | 2.61 | 0.35 |
| *Compound 6* | N/A | N/A | N/A | N/A | N/A | N/A | N/A |
| *Compound 7* | 21.35 | 9.72 | 20.78 | 0.45 | 0.97 | 11.81 | 3.51 |
| *Compound 8* | 85.95 | 31.57 | ≥ 60 | ≥ 60 | N/A | 22.73 | 0.00 |
| *Compound 9* | 172.27 | 172.11 | 0.20 | 0.03 | 0.00 | 0.24 | 0.23 |
| *Compound 10* | 10.66 | 0.90 | 17.80 | 8.51 | 1.67 | 8.27 | 5.46 |
| *Compound 11* | N/A | N/A | N/A | N/A | N/A | N/A | N/A |
| *Compound 12* | 5.17 | 0.32 | 13.55 | 0.89 | 2.62 | 7.73 | 2.67 |
| *Compound 13* | 7.27 | 3.26 | 10.66 | 3.24 | 1.47 | 13.84 | 3.88 |
| *Compound 14* | 7.69 | 0.63 | 15.30 | 1.78 | 1.99 | 11.82 | 1.48 |
| *Compound 15* | 4.32 | 0.00 | 10.51 | 0.00 | 2.43 | 2.43 | 0.00 |
| *Compound 16* | N/A | N/A | N/A | N/A | N/A | N/A | N/A |
| *Compound 17* | 19.90 | 1.61 | ≥ 60 | N/A | ≥ 3.01 | ≥60.00 | N/A |
| *Mocetinostat* | 4.79 | 0.16 | 6.39 | 0.51 | 1.33 | 4.88 | 0.03 |
| *Entinostat* | 38.32 | 2.94 | 20.80 | 0.69 | 0.54 | 32.27 | 2.69 |
| *PCI-34051* | 3.93 | 1.45 | 22.01 | 9.67 | 5.61 | 27.92 | 7.89 |
| *Tubastatin-A* | 2.03 | 1.23 | 3.58 | 1.16 | 1.76 | 3.40 | 0.44 |
| *MC-1568* | 1.10 | 0.27 | 10.68 | 8.23 | 9.69 | 18.58 | 2.53 |

